# SUITPy: A Python-based toolbox for the analysis of cerebellar functional and anatomical imaging data across the human lifespan

**DOI:** 10.64898/2026.05.14.724397

**Authors:** Yaping Wang, Yao Li, Bassel Arafat, Vahid Ashkanichenarlogh, Caroline Nettekoven, Ana Luísa Pinho, Carlos R. Hernandez-Castillo, Andre F. Marquand, Jörn Diedrichsen

## Abstract

The human cerebellum plays a central role in motor, emotional, and cognitive functions, and is implicated in many brain disorders. To improve the analysis of functional and anatomical imaging from the cerebellum, we introduce SUITPy, an improved and fully revised Python implementation of the widely used SUIT toolbox. For this new version, we developed a U-Net based model to automatically isolate the cerebellum from adjacent cortical tissue, which achieves higher fidelity than existing algorithms. The isolation works robustly without manual corrections for imaging data across the lifespan. We show that isolation and subsequent normalization to a cerebellum-only template lead to a more precise alignment of cerebellar structures across participants compared to normalization using a whole-brain template. We also show the utility of the cerebellar mask to prevent contamination of cerebellar functional data from surrounding cortical structures. The toolbox also provides functionality for visualizing cerebellar data on a flatmap, along with a range of anatomical and functional cerebellar atlases, thereby offering an essential tool that enables accurate cerebellar analysis across the lifespan.

## INTRODUCTION

Traditionally viewed as a motor structure, the human cerebellum is now recognized to support a broad range of cognitive, social, and emotional functions (Strick et al., 2009; Leiner et al., 1986; King et al., 2019; Nettekoven et al., 2024). Both anatomical and functional *Magnetic Resonance Imaging* (MRI) of the cerebellum play an increasingly important role in studies of neurodevelopmental, neurological, and psychiatric disorders (Schmahmann, 2004; Phillips et al., 2015). However, imaging the cerebellum remains challenging, as its surface is highly folded and tightly compacted within a small volume (Sereno et al., 2020). Furthermore, different regions of the human cerebellum subserve different functions, which are spatially located quite close together (Buckner et al., 2011; King et al., 2019; Nettekoven et al., 2024). Cerebellar cortical lesions therefore can have dramatically different consequences depending on their exact location.

To address this problem, we introduced a high-resolution cerebellum-only template—the *Spatially Unbiased Infratentorial Template* (SUIT)—together with a spatial normalization framework, to improve the inter-subject alignment of cerebellar structures (Diedrichsen, 2006). The analysis pipeline was shown at that time to outperform a common whole-brain normalization pipeline (Ashburner and Friston, 1999, implemented in SPM99). Since then, the SUIT toolbox has been widely adopted and used across a broad range of cerebellar imaging applications, including functional MRI (Baumann and Mattingley, 2012; King et al., 2019; Shahshahani et al., 2024), lesion (Timmann et al., 2008) and voxel-based morphometric studies (Moberget et al., 2018; Kim et al., 2025b). The SUIT pipeline employs a continuous voxel-wise approach, providing higher-resolution characterization than traditional lobular parcellation methods (Han et al., 2020; Faber et al., 2022). This is especially important because cerebellar functional organization does not align with anatomical lobular boundaries (King et al., 2019).

Despite its methodological advantages, the cerebellar isolation algorithm in the original SUIT toolbox does not always work robustly. For example, voxels belonging to the abutting neocortex or the transverse sinus are frequently misclassified as cerebellar tissue. In turn, inferior cerebellar regions are sometimes not included in the mask. This poses challenges for accurate anatomical analysis, as the volume of cerebellar tissue would not be accurately estimated. The incorrect isolation also biases the normalization to the cerebellum-only template, which propagates errors to subsequent analyses. The isolation mask therefore requires manual correction, limiting its applicability to large population-based MRI studies. Importantly, SUIT was originally developed using only data from healthy young adults, the algorithm performs poorly on developmental, aging, and clinical populations, where cerebellar anatomy deviates substantially from the normative template.

To address these challenges, we developed SUITPy, a Python toolbox for cerebellar MRI analysis across the lifespan. In this paper, we present a new deep learning–based cerebellar isolation algorithm using a U-Net architecture (Ronneberger et al., 2015) that works robustly across different age groups and imaging resolutions, and can leverage either T1- or T2-weighted images or both when available. We show that using this isolation mask during the normalization to a cerebellum-only template outperforms current state-of-the-art whole-brain normalization approaches. The cerebellar isolation also prevents the intrusion of cortical signals into the cerebellum in functional MRI analyses. The toolbox additionally includes tools for surface-based visualization and a collection of cerebellar atlases, providing a comprehensive and accessible resource for the neuroimaging community. Taken together, SUITPy provides a robust framework for cerebellar anatomical and functional MRI analysis across the human lifespan.

## METHODS

### Datasets

To ensure coverage of the entire lifespan, we compiled a collection of anatomical images from eight MRI datasets with varying image resolution, contrasts, and quality (n=131; age 0–76 years, see Table 1). The dataset included both publicly available datasets and datasets that were acquired at Western University and at the University of Essen. All procedures for the locally acquired datasets were approved by the relevant institutional ethics boards, and data transfer agreements were established for the reanalysis of anonymised data.

**Table 1.** Dataset for U-Net isolation algorithm development. *dHCP*: Developing Human Connectome Project (Edwards et al., 2022). *BCP*: The Baby Connectome Project (Howell et al., 2019). *HCP-D* Child and Adolescent Data from the Human Connectome Project – Development (Somerville et al., 2018). *MDTB*: Multi Domain Task Battery (King et al., 2019). *HResMDTB*: High resolution MDTB (unpublished data). *TDTB*: Timing Domain Task Battery (https://github.com/alpinho/tdtb_analysis). These three datasets were collected at Western University. *HCP-Y*: Human Connectome Project - young adult (Van Essen et al., 2012). *EssCer*: dataset comprising patients with cerebellar degeneration and age-matched controls collected at the University of Essen (Draganova et al., 2022).

| Source | Field strength | T1w resolution | N | Age/Years<br>Mean±SD [range] | Sex<br>M/F | T1w | T2w |
| --- | --- | --- | --- | --- | --- | --- | --- |
| dHCP | 3T | 0.5mm | 30 | 0.02 ± 0.03 [0.00, 0.09] | 16/14 | Yes | Yes |
| BCP | 3T | 0.8mm | 10 | 2.09 ± 0.77 [1.08, 3.17] | 5/5 | Yes | Yes |
| HCP-D | 3T | 0.8mm | 10 | 16.60 ± 3.53 [10, 21] | 6/4 | Yes | Yes |
| MDTB | 3T | 1mm | 24 | 23.29 ± 2.82 [19, 29] | 8/16 | Yes | No |
| HResMDTB | 7T | 0.7mm | 7 | 25.00 ± 3.27 [21, 29] | 3/4 | Yes | Yes |
| HCP-Y | 3T | 0.7mm | 10 | 29.10 ± 3.78 [23, 33] | 5/5 | Yes | Yes |
| TDTB | 3T | 1mm | 20 | 25.80 ± 6.79 [18, 41] | 10/10 | Yes | No |
| EssCer | 3T | 0.5mm | 20 | 59.15 ± 11.13 [34, 76] | 10/10 | Yes | No |

Specifically, for the neonatal cohort, we included 30 subjects from the Developing Human Connectome Project (dHCP) (Edwards et al., 2022), with postnatal ages of 0–4 weeks. The infant subjects (n=10, age 0–5 years) were randomly sampled from the Baby Connectome Project (BCP) (Howell et al., 2019). The child and adolescent subjects (n=10, age 10–21 years) were randomly selected from the Human Connectome Project – Development (HCP-D) (Somerville et al., 2018). The young adult MRI scans (n=61) were aggregated from multiple independent collections, including the HCP Young Adulthood (HCP-Y) (Van Essen et al., 2012), the Multi-Domain Task Battery (MDTB) (King et al., 2019), the high resolution MDTB (HResMDTB, unpublished data), and the Timing Domain Task Battery (TDTB) datasets. To represent elderly populations and pathological conditions, we included 10 controls and 10 patients from a dataset comprising patients with cerebellar degeneration and age-matched controls acquired at the University of Essen (EssCer) (Draganova et al., 2022).

Across different datasets, we therefore captured broad demographic and neuroanatomical variability. The dataset also differed in imaging hardware (3T and 7T scanners), MRI quality, contrast and resolution. This cross-site heterogeneity supports robust model generalization.

### Cerebellar manual isolation

For the resulting N=131 individual scans, a total of 6 expert raters manually generated cerebellar masks. The raters started with an initial mask generated by the probabilistic isolation algorithm from the SUIT toolbox (Diedrichsen, 2006). The masks were discretized at a probability threshold of p=0.2 and then manually corrected. Each mask was corrected by one rater using FSLeyes (McCarthy, 2025) by going through all slices in axial, sagittal, and coronal orientation. The mask was then reviewed by a second rater, and any discrepancies were discussed between the two raters and resolved by consensus.

The goal was to generate an isolation mask that included cerebellar white and gray matter, as well as the brainstem. The reason for including the brainstem is twofold: first, the brainstem includes important structures (inferior olive, pontine nuclei) to understand the cerebellar system and is therefore often of interest. Secondly, by including the brainstem, we can avoid the difficulty of defining the boundary between cerebellum and brainstem, which would require drawing a somewhat arbitrary plane through the middle cerebellar peduncle. Uncertainty about this boundary would influence the subsequent normalization algorithm or add variability to the subsequent white-matter volume estimates. We therefore included the brainstem from the medulla oblongata inferiorly to the midbrain superiorly. This ensured that the normalization of the cerebellum and the brainstem structures could be clearly interpreted.

### Cerebellar probabilistic isolation

As a baseline, we used the isolation algorithm implemented in the previous Matlab-based version of SUIT (Diedrichsen, 2006). This algorithm is based on SPM’s unified segmentation and normalization algorithm (Ashburner and Friston, 2005). In the earliest version of SUIT, the boundary between cerebellar and neocortical gray matter was determined using the distance of each voxel in question from the cerebellar and neocortical white-matter body. With the release of version `3.1`, we implemented a newer version of the algorithm that introduced separate tissue priors for cerebellar and cortical gray and white matter (`suit_isolate_seg.m`). We used this improved, newer version throughout the paper.

### Cerebellar U-Net Isolation

#### Preprocessing

The model was trained to produce a cerebellar isolation mask including all cerebellar gray matter and white matter, as well as the brainstem. The input to the model was either a T1-weighted or T2-weighted image (or both). To reduce the computational load on training and to ensure that the algorithm would work well on different brain sizes and images of different resolutions, we first cropped the images to a fixed 128 × 128 × 128 voxel bounding box centered on the cerebellum and brainstem.

This was achieved by aligning the input image to the `MNI152NLin6Asym` template (Grabner et al., 2006) using an affine registration. This transformation, comprising uniform scaling, rotation, and translation, establishes a consistent cerebellar–brainstem coordinate frame for subsequent region definition, while preserving the unique anatomical characteristics of each subject for the model to learn.

By default, the algorithm used the T1-weighted image (or the T2-weighted image if the T1-weighted image was not available) to compute affine registration with the template. To ensure robust affine registration, the algorithm computed the normalized mutual information (Studholme et al., 1999) between the original image and the template. Based on visual inspection, a normalized mutual information value above 1.23 indicates decent alignment with the template. When the value falls below this threshold, the algorithm iteratively repeated the registration from different starting values to achieve a better alignment. Once the transformation matrix was calculated, the input image was resliced into the template space and cropped.

#### U-Net architecture

Recent research has demonstrated the advantages of U-Net style architectures in the segmentation of volumetric medical images (Ronneberger et al., 2015; Zhou et al., 2018; Huang et al., 2020; Isensee et al., 2018; Hatamizadeh et al., 2022), showing that U-Net and its variants are competitive due to their strong inductive bias for spatial localization and their ability to perform well even with limited training data. Compared to purely transformer-based models, U-Net provides a better trade-off between computational efficiency and segmentation accuracy in high-resolution 3D tasks.

The U-Net architecture comprises a symmetric encoder–decoder design and skip connections. The symmetric encoder-decoder structure plays a critical role in feature learning. The encoder progressively reduces spatial resolution while increasing feature abstraction, enabling the network not only to focus on local features but also to capture global contextual information such as the overall cerebellar shape. Thus, the hierarchical encoder extracts multi-scale features, which are essential for capturing both the overall cerebellar shape (global feature) and fine structural boundaries (local details). Multi-scale representation is particularly important in medical imaging, where structures of interest vary significantly in size and appearance (Hatamizadeh et al., 2022). The decoder restores spatial resolution through progressive up-sampling combined with skip connections, enabling precise localization, which is critical for accurately delineating the boundaries between the cerebellum and adjacent brain structures. This symmetry ensures that coarse global information and fine spatial details are effectively integrated (Çiçek et al., 2016; Hatamizadeh et al., 2022). Moreover, skip connections are a key component that directly addresses the loss of spatial information during down-sampling. By transferring high-resolution features from the encoder to the decoder, they preserve boundary and edge information that would otherwise be lost. Additionally, skip connections improve gradient flow during training and stabilize optimization, leading to better convergence and performance (Chenarlogh et al., 2024).

For this project, we implemented a 3D U-Net for volumetric MRI data (Fig. 1). The model took two channels of 128 × 128 × 128 volume containing co-registered T1-weighted and T2-weighted images as input. If one of the input images is not present, all voxels for that channel were set to zero. The use of dual-channel input allows the network to leverage complementary contrast information from different MRI modalities. This multi-modal fusion has been shown to improve segmentation accuracy in brain imaging tasks (Hatamizadeh et al., 2022). Setting missing modalities to zero enables the model to handle incomplete data without requiring architectural changes.

**Figure 1.**
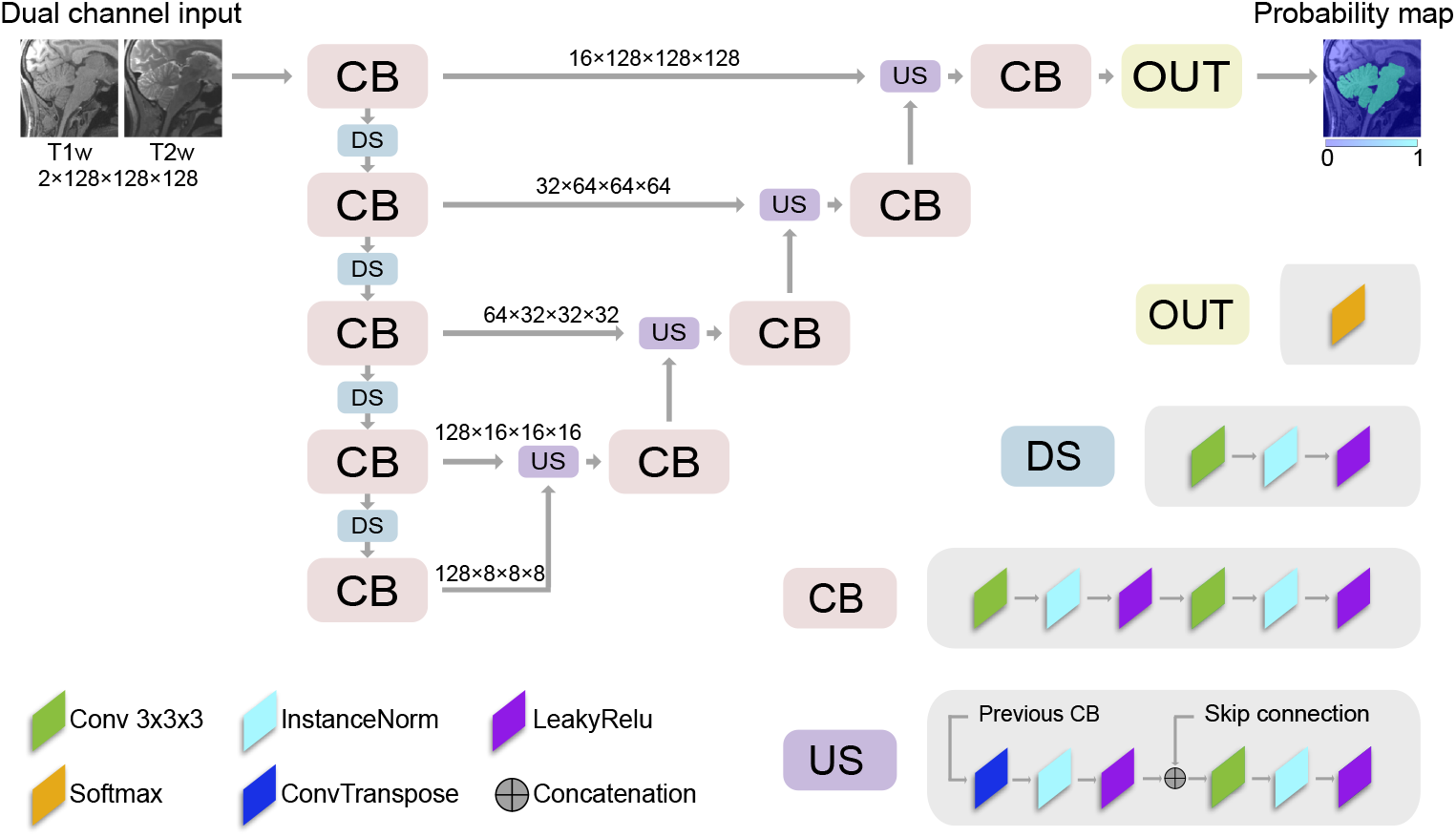
Architecture of U-Net for cerebellar isolation. The network follows a symmetric encoder-decoder architecture with skip connections. The encoder progressively compresses spatial resolution through a series of Convolutional Blocks (**CB**) and Down-Sampling Blocks (**DS**), while the decoder reconstructs the spatial resolution through Up-Sampling Blocks (**US**) and CBs. Each CB consists of two repeated sequences of a 3D convolutional layer, InstanceNorm, and LeakyReLU. Each DS applies a strided convolution followed by InstanceNorm and LeakyReLU. Each US applies a transposed convolution followed by InstanceNorm and LeakyReLU, and concatenates the result with the corresponding skip connection from the encoder. The concatenated data goes through another sequence of 3D convolutional layer, InstanceNorm, and LeakyReLU before passing it to the next CB. The Output Block (**OUT**) applies a softmax activation to produce the final probability map.

The network was composed of four sequential encoding and decoding stages. The first encoding stage used 16 feature channels, with subsequent stages increasing to 32, 64, and 128 channels, respectively. The depth of four encoding stages was selected to balance representational capacity and computational efficiency. Fewer stages did not provide sufficient receptive field expansion to capture the full cerebellar context. Deeper architectures incurred unnecessary computational cost without demonstrable performance gains.

Each encoding stage started with a Convolutional Block (CB), which consisted of two sequential non-linear combinations. Each combination started with a 3D convolution layer with a kernel size of 3, a stride of 1, and appropriate padding to preserve the spatial dimensions of the feature maps. This was followed by an Instance Normalization with affine set to true across the entire image (Ulyanov et al., 2016) and a leakyReLU with a negative slope of 0.01 as a non-linearity (Maas, 2013). LeakyReLU enhances gradient flow by introducing a small, non-zero slope in the negative region, improving training stability (Sayın et al., 2025). Applying this combination twice increases CB’s capacity and receptive field within the block.

After each CB, there was a Down-Sampling Block (DS), which again consisted of a convolution, instance normalization, and leakyReLU. In contrast to the CB, the convolution was applied with a stride of 2, thereby reducing the overall image resolution. Using strided convolution rather than max-pooling enables the DS to learn the downsampling operation, thereby increasing the model’s capacity.

After the final DS, the feature maps passed through the CB at the bottom of the network, referred to as the bottleneck. This represents the most compressed latent space representation of the input, where all high-level features are abstracted before being passed to the decoding path.

Each of the four decoding stages started with an Up-Sampling Block (US), followed by a CB. The US not only performed up-sampling on the data from the previous block, but also received data from the skip connection and concatenated them. Within the US, the data from the previous CB was passed through a transposed convolution with a 2×2×2 kernel and a stride of 2, followed by InstanceNorm and LeakyReLU, allowing the network to learn spatial resolution reconstruction rather than relying on fixed interpolation. The resulting feature maps were then concatenated with the corresponding skip connection from the encoder CB, to preserve full feature information from both encoder and decoder pathways, to increase representational capacity and improve precision (Çiçek et al., 2016). The resulting features from the US were then refined by the subsequent CB before being passed to the next up-sampling stage.

The network concluded with a final Output Block (OUT) that mapped features to probability maps via a softmax operation. Thus, the network’s direct output was a voxel-wise probability map of the cerebellum–brainstem mask in the 128 × 128 × 128 field of view.

#### Model training

We formulated cerebellar isolation as a voxel-wise binary classification task, where each voxel was predicted as either cerebellum–brainstem tissue or background. We therefore chose Cross Entropy loss as the objective function, as it is suitable for probabilistic classification problems. The Adam optimizer (Kingma and Ba, 2017) was used with default parameters. The model was trained for 200 epochs, at which point the loss had converged and test performance plateaued.

To make the best use of the data and improve the model’s robustness across different input modality combinations, we designed a specialized training strategy. The training aimed to achieve two objectives. First, the model should be able to detect zero-padding channels (a missing T1 or T2 image) and learn to ignore them. Second, when both channels are provided, the model should learn the connections between the two modalities and integrate complementary features to enhance prediction accuracy.

To achieve these goals, from each subject for whom both T1w and T2w images were available, we created three separate training samples: (1) T1w image with zero padding, (2) T2w image with zero padding, and (3) T1w and T2w images jointly. These samples were combined with the subjects that had only a single modality image to form a data set of 131 T1w samples, 67 T2w samples, and 67 joint T1w and T2w samples.

To supplement the limited dataset size and to improve the model’s generalizability across different cerebellar locations and orientations, we also performed geometric data augmentation. For each original image(s), we introduced random rotation and shifting after the image was transformed into template space. The images were rotated along three axes, with rotation angles sampled from a normal distribution with a mean of 0^◦^ and a standard deviation of 15^◦^. After rotation, the images were shifted along the three axes. Shift distances in the X and Y directions were sampled from a normal distribution with a mean of 0 mm and a standard deviation of 3 mm, while the shift distance in the Z direction was sampled from a normal distribution with a mean of 0 mm and a standard deviation of 5 mm. For each image, we created three sets of augmented data in the training set, resulting in a total training set four times larger than the original.

The model ultimately implemented in the SUITPy toolbox was trained using all 131 subjects and all available modality combinations, yielding 265 data samples. The geometric data augmentation techniques were applied to generate an additional 795 samples, resulting in a total training dataset of 1060 samples.

#### Postprocessing

Because model inference is performed in template space, the predicted probability maps needed to be resampled into the individual native anatomical space by applying the inverse of the initial affine transformation used during registration of the preprocessing, ensuring accurate alignment with the original scan.

These resampled probability maps were then binarized using a 0.5 threshold to generate final isolation masks. Finally, to eliminate any spurious voxels, a connected-component analysis was performed on the binary mask, retaining only the largest contiguous cluster.

#### Evaluation of U-Net mask

We evaluated the performance of the U-Net by comparing the predicted mask to the manually corrected ground-truth masks. To avoid bias from different image resolutions, this evaluation was performed in the standardized 128 × 128 × 128 voxel bounding box in the template space. We employed two complementary quality metrics. First, we used the Hausdorff Distance (Huttenlocher et al., 1993): For each voxel in image A, we found the closest voxel in image B, and then found the maximum of these distances across the entire image. The procedure was then repeated, switching the role of the two images. The maximum of these two numbers was taken as a symmetric distance measure. Overall, the Hausdorff distance therefore found the place where the predicted boundary deviated the furthest from the ground truth boundary.

Additionally, we used the dice score (Sørensen, 1948; Dice, 1945) to measure the volumetric overlap between the predicted and the ground truth mask. A dice score of 1 indicates perfect overlap, while 0 indicates no overlap. While the dice score provides a robust measure of overall accuracy, it mostly reflects the overlap of the volumes and is less sensitive to subtle boundary errors than the Hausdorff distance. Because such boundary voxels constitute a small fraction of the total 3D volume, their misalignment may not substantially reduce the dice score, even when clinically relevant.

Our ground-truth masks included the brainstem to avoid any arbitrary boundaries between the cerebellar white matter and brainstem. However, the inferior (below the medulla oblongata) and superior (above the mesencephalon) boundaries of the mask were somewhat arbitrary. Consequently, we did not expect that the model would predict these boundaries well. Therefore, we excluded the upper and lower 5mm of the brainstem from the evaluation by defining an evaluation mask in the bounding box.

To evaluate model performance, we used a four-fold cross-validation scheme. Each dataset was randomly partitioned into four non-overlapping folds. In each iteration, three folds (≈ 75% of the data) served as a training set, and the remaining fold (≈ 25%) was held out for testing. The process was repeated four times so that every scan was used as an independent test sample exactly once. The regime ensured that both training and testing sets included the full breadth of datasets.

We compared our new model with the probabilistic isolation implemented in SUIT (Diedrichsen, 2006). Additionally, we assessed the extent to which affine alignment to the template space alone, without any learning-based refinement, would already superimpose cerebellar structures across subjects. To this end, we averaged the manual isolation masks from the training set in the 128 × 128 × 128 voxel bounding box after affine alignment. The resulting mean mask was then discretized (threshold = 0.5) and used as a baseline prediction for the test set.

### Cerebellar normalization

Non-linear normalization into a group template system allows for group-level analyses at the voxel level. For anatomical analysis, the non-linear normalization also provides a metric of the relative size of each part of the cerebellum in relationship to the template.

The original SUIT toolbox employed the DARTEL algorithm (Ashburner, 2007) to normalize the images to a cerebellum-only template. We showed that, compared with the conventional whole-brain normalization using basis functions (Ashburner and Friston, 1999), this approach resulted in a 10% increase in group-level statistical t-values within activated regions (Diedrichsen, 2006). For the Python implementation, we used a comparable diffeomorphic algorithm implemented in the ANTs toolbox through `antsRegistrationSyN` (Avants et al., 2011). To ensure a direct comparison between whole-brain and cerebellum-only approaches, we generated a new cerebellum-only template by masking the cerebellum and brainstem from the symmetric `MNI152NLin2009cSym` template (Fonov et al., 2009, 2011), i.e., the `MNI152NLin2009cSymC` template. We used the output of the U-Net isolation to mask the original images in native space and then aligned them to the cerebellum-only template.

We then assessed whether this cerebellum-only pipeline improved spatial normalization compared to two standard imaging pipelines. Both pipelines started with an N4 bias-correction (Tustison et al., 2010) of the anatomical image. In the whole-head pipeline, we normalized the entire T1-weighted image to the `MNI152NLin2009cSym` template. In the whole-brain pipeline, we skull-stripped (Avants et al., 2011) the original image and then registered it to a skull-stripped version of the `MNI152NLin2009cSym` template.

### Evaluation of normalization

To evaluate the performance of different normalization pipelines, we used two approaches. First we calculated the correlation coefficient between the template and the resliced image. To ensure a fair comparison, we defined an evaluation mask that encompassed the cerebellar-only template area, including cerebellar gray matter, white matter, the pons, and the medulla oblongata, while excluding the brainstem and midbrain.

In order to test how well each pipeline aligned the internal structures of the cerebellum across individuals, we used ACAPULCO (Han et al., 2020) to obtain a lobular parcellation on each individual. These lobular masks were resliced into group space. The generalized dice score (Crum et al., 2006) was calculated for each pair of subjects within the same dataset, yielding a subject × subject similarity matrix. For each subject, we averaged the dice score with all other subjects. This provided a measure of how consistently the lobular parcellation for a given subject aligns with those of the remaining cohort.

For the statistical analysis, we assessed normalization performance at both the subject and dataset levels. At the subject level, we applied paired t-tests to compare correlation or dice score across pipelines, evaluating whether normalization quality differed significantly across methods for the same individuals. At the dataset level, we first computed the average correlation or dice score across all subjects within each dataset, then performed paired t-tests using each dataset as an independent sample.

### Functional MRI analysis

To quantify the additional impact of applying the cerebellar isolation mask during functional data reslicing — beyond the benefits already conferred by cerebellum-only normalization — we used anatomical and functional data from two datasets: the Multi-Domain Task Battery (MDTB; N=24, functional resolution 2.5 × 2.5 × 3.0 mm^3^) (King et al., 2019) and the Language multi-task battery (N=17, functional resolution 2.3 × 2.3 × 2.3 mm^3^) (Arafat et al., 2026).

For each subject, the anatomical T1-weighted image was processed using the U-Net based isolation without subsequent manual correction, followed by normalization to the cerebellum-only template, i.e., SUIT space. We also segmented the anatomical T1-weighted image into white matter, gray matter, and cerebrospinal fluid using the unified segmentation algorithm implemented in SPM (Ashburner and Friston, 2005). To simulate the degree to which fMRI contrast maps within cerebellar voxels reflect neocortical versus cerebellar activity, we resampled these anatomical maps to the space of the individual fMRI space acquisition. Subsequently, these maps were resliced into SUIT atlas space, either with or without application of the cerebellar isolation mask. All resliced images were then spatially smoothed across all non-zero values in the image using AFNI’s `3dBlurToFWHM` (Cox, 1996) with Gaussian kernels of varying width (2, 4, 6, 8, and 10 mm FWHM). Then the proportion of functional signal arising from the neocortex was estimated by dividing the cortical gray and white matter probability by the sum of cortical and cerebellar gray and white matter probability. These normalized cortical probability maps were averaged across subjects to obtain a group-average map, which was visualized using the surface-based flatmap representation of the cerebellar cortex implemented in SUITPy (Diedrichsen and Zotow, 2015).

## RESULTS

### Cerebellar isolation comparison between U-Net and probabilistic isolation algorithm

The probabilistic isolation algorithm of the legacy SUIT toolbox tends to have a number of characteristic problems and shortcomings in correctly delineating the cerebellar boundary. It frequently misclassifies adjacent structures, such as the part of the inferior occipital-temporal cortex (Fig. 2c), the transverse sinus (Fig. 2d), or fails to include parts of cerebellar tissue (Fig. 2e). More critically, the probabilistic isolation algorithm often fails dramatically in populations other than healthy young controls, such as neonates (Fig. 2f) or individuals with cerebellar degeneration (Fig. 2g).

**Figure 2.**
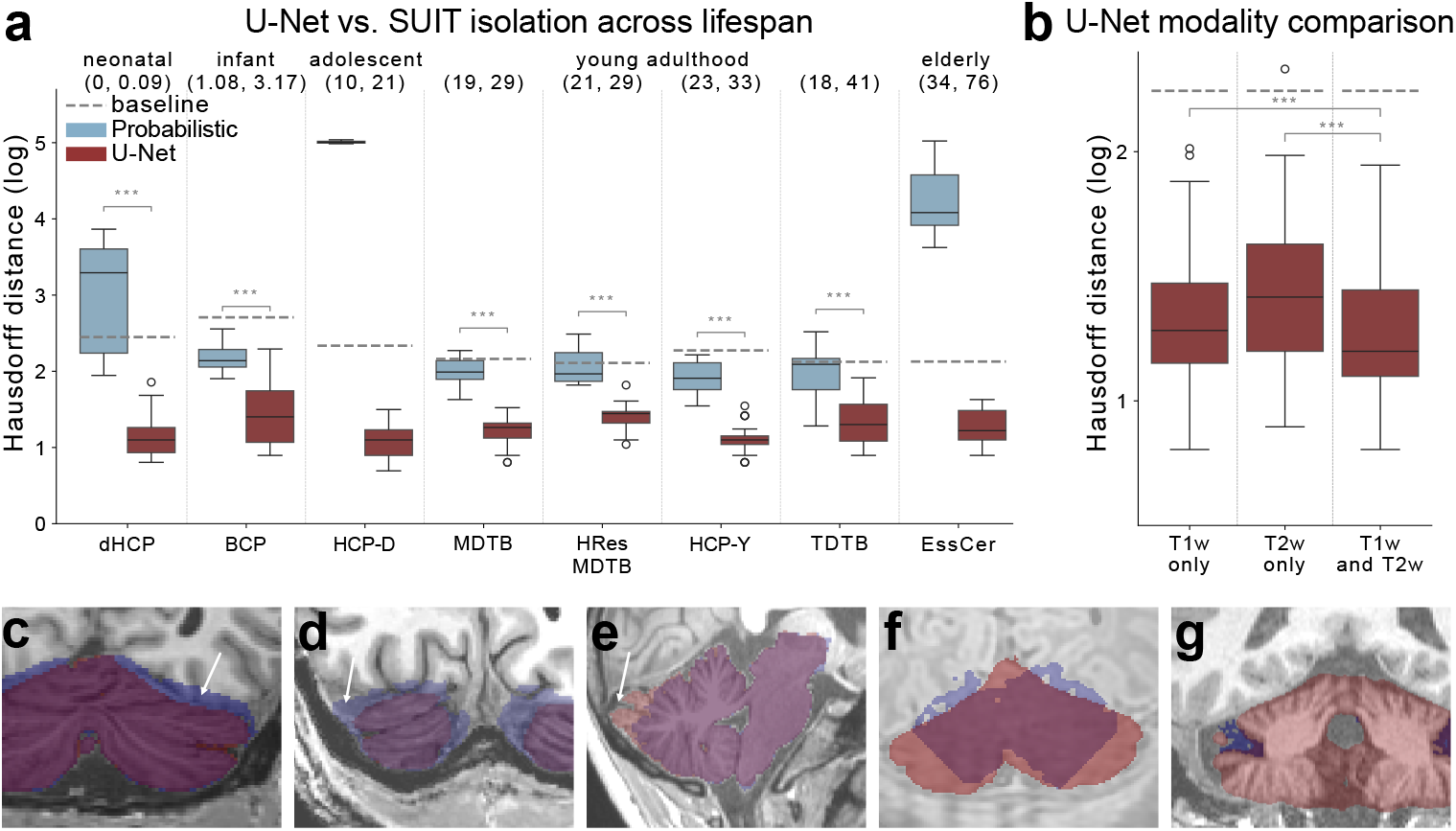
Comparison of U-Net and probabilistic isolation algorithm. **(a)** Hausdorff Distance (log scale) for probabilistic (blue) and U-Net (red) isolation. Dotted lines indicate baseline performance of the simple average of the training set. Horizontal bars with asterisks denote significant pairwise differences (paired t-tests, *: p<0.05, **: p<0.01, ***: p<0.001). **(b)** Hausdorff Distance (log scale) of U-Net isolation using different input modality configurations. **(c–g)** Example masks comparing the legacy probabilistic isolation (blue) and U-Net isolation (red): **(c)** probabilistic mask incorrectly includes occipital lobe gray matter; **(d)** probabilistic mask incorrectly includes transverse sinuses; **(e)** midsagittal vermis excluded from probabilistic mask; **(f)** near-complete failure of probabilistic mask in a neonatal subject; **(g)** complete failure of probabilistic mask in an elderly subject. In contrast, the U-Net model produces adequate masks across all cases.

To improve the isolation of the cerebellum from surrounding tissues, we trained a U-Net (Ronneberger et al., 2015) deep learning model on manual isolation data from 8 datasets, which spanned the entire lifespan (0–76 years), utilized different imaging resolutions and modalities (T1- or T2-weighted images), and also included cerebellar degeneration patients (Table 1).

We first compared our proposed U-Net model against the probabilistic isolation algorithm on young healthy adult datasets, where the legacy probabilistic isolation was known to perform most reliably. In all adult datasets, including MDTB, HResMDTB, HCP-YoungAdult, and TDTB, the U-Net isolation consistently achieved lower Hausdorff Distance (Huttenlocher et al., 1993) than the probabilistic isolation (Fig. 2a, all *p <* 1.316 × 10^−6^). Similar results were achieved when using the Dice score (Fig. S1).

More importantly, the U-Net isolation also performed well on datasets for which the probabilistic isolation historically failed, for example, the neonatal data from dHCP (ages 0–0.09 years) (Fig. 2f) and the elderly subjects with or without cerebellar degeneration (Fig. 2 g). For these datasets, the probabilistic parcellation performed worse than the baseline of using a group-average mask after affine normalization (see Evaluation of U-Net mask for details). Across the entire dataset, the legacy probabilistic isolation algorithm failed altogether for 24.4% of the cases, with most failures occurring in neonates and elderly subjects. In contrast, the new U-Net model resulted in robust isolation across all datasets and yielded acceptable performance on all subjects from all datasets, each time clearly outperforming baseline performance.

### Isolation performance using different imaging modalities

In clinical and research settings, datasets frequently contain images of varying modalities. As T1w and T2w images provide complementary structural information, an ideal model should perform reliably on either single-modality inputs while potentially improving when both are available. To this end, our U-Net was designed to have two input channels, and we optimized the training strategy to accommodate missing modalities (see Model training for details). To assess the benefit of multiple input modalities, we organized a dataset of subjects for whom both T1- and T2-weighted scans were available and trained and evaluated the model’s performance on it. We compared the performance when only the T1w was given, when only the T2w was given, and when both were provided.

The comparison (Fig. 2b) showed that the U-Net model performed robustly with T1w or T2w alone. Performance in both modalities was substantially better than both probabilistic isolation and the baseline. When both T1w and T2w images were provided, the model consistently yielded more accurate masks, outperforming T1w alone (*t*_36_ = 3.660, *p* = 8.030 × 10^−4^) as well as T2w alone (*t*_36_ = 3.663, *p* = 7.951 × 10^−4^), although the magnitude of this improvement was modest. This suggests that the U-Net model can leverage complementary contrast information from both modalities, although the practical benefit of providing both may be limited.

### Cerebellar isolation improves spatial normalization

Diedrichsen (2006) demonstrated a substantial improvement in cerebellum-only normalization accuracy compared to what was the state-of-the-art whole-brain pipeline at the time (Ashburner and Friston, 1999). Since then, both normalization algorithms (e.g., Ashburner, 2007; Avants et al., 2011) and whole-brain templates (Grabner et al., 2006; Fonov et al., 2009, 2011) have improved considerably. Therefore, it was unclear whether cerebellum-only normalization still conferred an advantage when compared to equivalent whole-brain approaches. We therefore compared the cerebellum-only pipeline (using the U-Net derived cerebellar mask) to a whole-head pipeline to the MNI2009cSym template (Fonov et al., 2011) and whole-brain pipeline, in which the skull-stripping (Avants et al., 2011) was employed before normalization.

We first evaluated the quality of the normalization by correlating the deformed individual images with the template. To ensure a fair comparison, the correlation was only calculated within the cerebellum and brainstem (Fig. 3c), for which the two templates (MNI2009cSym and cerebellum-only MNI2009cSymC) were identical.

**Figure 3.**
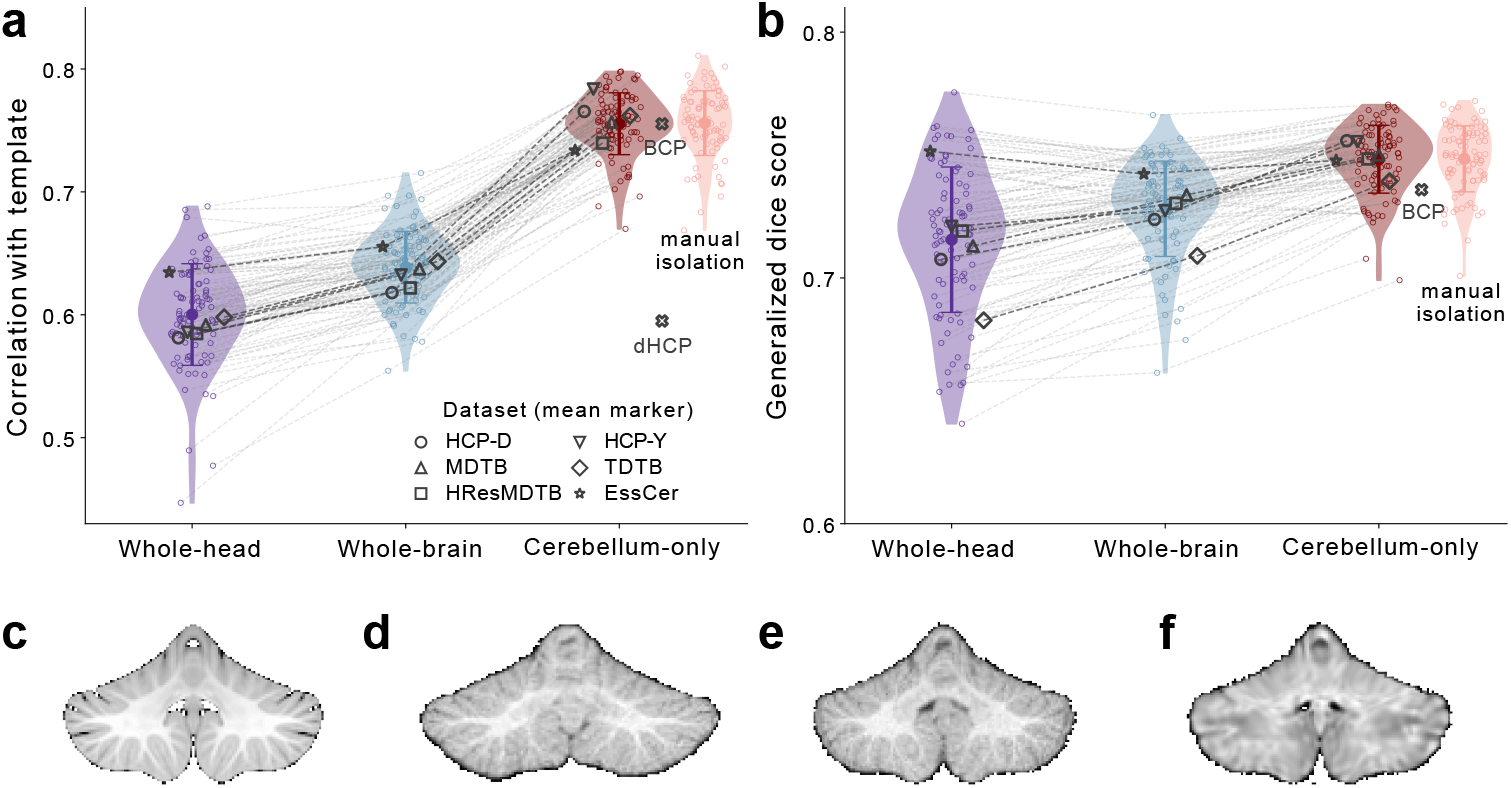
Cerebellar isolation improves spatial normalization. **(a)** Pearson correlation between the MNI2009cSymC template and the individual image after normalization with whole-head (purple), whole-brain (blue), or cerebellum-only pipeline (red). Performance of the cerebellum-only pipeline using the manual isolation mask (pink) is shown for comparison. Correlation is computed across cerebellar voxels, as defined by the template mask shown in **(c)**. Each dot represents an individual subject; larger symbols are the mean for each dataset. For the neonatal (dHCP) and infant (BCP) datasets (cross symbol), both the whole-brain and whole-head pipelines failed, so only the cerebellum-only pipeline is shown. **(b)** Lobular inter-subject alignment after normalization. Each dot shows the average generalized dice score for each subject with all other subjects within the same dataset. Larger symbols denote dataset-level means. For the infant dataset (BCP, cross symbol), both the whole-brain and whole-head pipelines failed, so only the cerebellum-only pipeline is shown. For dHCP the automated lobular segmentation with ACAPULCO failed. **(c)** The cerebellum-only MNI2009cSymC template. **(d–f)** Example normalized brains across different pipelines for infant and neonatal brains: **(d)** the whole-head pipeline provides suboptimal results for an example infant brain (BCP); **(e)** the cerebellum-only pipeline performs better for the same infant brain (BCP); **(f)** the cerebellum-only pipeline also produces accurate normalization for the neonatal brain (dHCP) despite image contrast inversion.

Across all individuals from the adolescents, young adults, and older adults datasets (Fig. 3a), the whole-brain pipeline outperformed the whole-head pipeline significantly (*t*_90_ = 13.167, *p* = 1.064 × 10^−22^), showing the advantage of skull-stripping before normalization (Avants et al., 2011). For these individuals, the cerebellum-only pipeline significantly outperformed both the whole-head (*t*_90_ = 33.603, *p* = 1.020 × 10^−52^) and whole-brain (*t*_90_ = 37.188, *p* = 2.041 × 10^−56^) pipelines. This advantage was present for each of these datasets. Even if we considered the dataset to be our random effect (i.e., averaging the correlation within each dataset and then testing across datasets), the cerebellum-only pipeline significantly outperformed the whole-head (*t*_5_ = 11.587, *p* = 8.404 × 10^−5^) and whole-brain pipeline (*t*_5_ = 11.418, *p* = 9.023 × 10^−5^).

In addition to voxel-based alignment to the template, we also asked whether the cerebellum-only pipeline would improve the superposition of cerebellar anatomical features across individuals. To do this, we obtained individual lobule parcellations using ACAPULCO (Han et al., 2020), and then resliced the individual lobular mask into template space using the transformation estimated from each pipeline. We calculated the average generalized dice score (Crum et al., 2006) for each participant with each of the other participants in the same dataset (Fig. 3b).

The results generally confirmed our previous analysis. Skull-stripping before normalization improved superposition of cerebellar structure, as can be seen in the significant difference between whole-head vs. whole-brain pipelines (*t*_90_ = 7.501, *p* = 4.304 × 10^−11^). The cerebellum-only pipeline again outperformed both the whole-head (*t*_90_ = 12.761, *p* = 6.741 × 10^−22^) and whole-brain pipelines (*t*_90_ = 14.736, *p* = 1.018 × 10^−25^). This advantage was also significant at the dataset level: cerebellum-only vs. whole-head pipelines (*t*_5_ = 3.960, *p* = 0.011), cerebellum-only vs. whole-brain pipelines (*t*_5_ = 5.171, *p* = 0.004).

Finally, we aimed to determine if the use of the fully automatically generated U-Net isolation mask was sufficient, or whether normalization could be improved even more using hand-corrected masks. We therefore repeated the cerebellum-only pipeline using our manual isolation rather than with the U-Net isolation (Fig. 3, pink). No significant difference was observed on correlation with template across individual level (*t*_90_ = 1.139, *p* = 0.258) and dataset level (*t*_5_ = 0.046, *p* = 0.965), as well as generalized dice score across individual level (*t*_90_ = 0.310, *p* = 0.757) and dataset level (*t*_5_ = 0.414, *p* = 0.696). This showed that the full advantage of a cerebellum-only normalization can be achieved with the new U-Net based normalization algorithm, without the need for manual intervention.

### Cerebellum-only normalization works robustly on infant and neonatal brains

The cerebellum-only pipeline also had the additional advantage of working for very young individuals. For the neonatal and infant brains (BCP and dHCP), the traditional whole-head and whole-brain normalization to the adult template nearly always failed (Fig. 3d) relative to the template (Fig. 3c). In contrast, restricting normalization to the cerebellum using a cerebellum-specific U-Net mask resulted in successful normalizations (Fig. 3e, f).

For infant brains (BCP), the cerebellum-only pipeline achieved template correlations comparable to adult datasets (*r* = 0.755 *±* 0.010, mean± standard deviation, Fig. 3a), as well as good inter-individual alignment of the lobules (dice = 0.736 *±* 0.009, Fig. 3b). Neonatal data (dHCP) exhibited contrast inversion in T1-weighted images, with white matter appearing darker than gray matter. Consequently, the correlations with template were substantially lower (*r* = 0.595 *±* 0.043, Fig. 3a, f). While the general alignment to the template remained acceptable, lobular evaluation was not feasible because ACAPULCO failed on neonatal brains.

### Cerebellar isolation prevents leakage of cortical signal in functional MRI analysis

While the cerebellum-only normalization improved alignment of cerebellar structures, we further investigated a second important use of the cerebellar isolation mask: preventing leakage of cortical signal into the cerebellum in functional MRI analysis. In conventional fMRI analysis pipelines, functional data are typically resliced into a whole-brain template and then spatially smoothed prior to group-level analysis. Because the anterior cerebellum is only separated by a thin membrane (the tentorium) from the inferior occipital cortex, this procedure can lead to systematic leakage of cortical signal into anterior cerebellar voxels. This effect is especially serious because the task-related fMRI signal is often substantially stronger in the neocortex than in the cerebellum.

To quantify the strength and spatial distribution of this signal leakage, we simulated the process of reslicing subject-specific fMRI maps into atlas space and then smoothed these with spatial kernels of varying sizes. Instead of real function maps, we used subject-specific cortical and cerebellar tissue gray and white matter maps to approximate what proportion of the fMRI signal in cerebellar atlas voxels would reflect real cerebellar vs. dislocated neocortical signal (see Functional MRI analysis).

Following standard fMRI group analysis, we observed substantial spurious cortical signals with a characteristic distribution in cerebellar lobules IV–VI (Fig. 4a). Averaged across the entire right lobule V, the mean cortical leakage reached 1.3% *±* 0.4% at 6mm smoothing. In the superficial voxels in the cerebellum, leakage was substantially higher, with some voxels reaching 21.1%. As expected, the degree of signal leakage increased monotonically with smoothing kernel size (Fig. 4b, dashed lines). The leakage effects were also more pronounced in fMRI datasets with lower resolution compared to higher resolution (3.0 mm vs. 2.3 mm).

**Figure 4.**
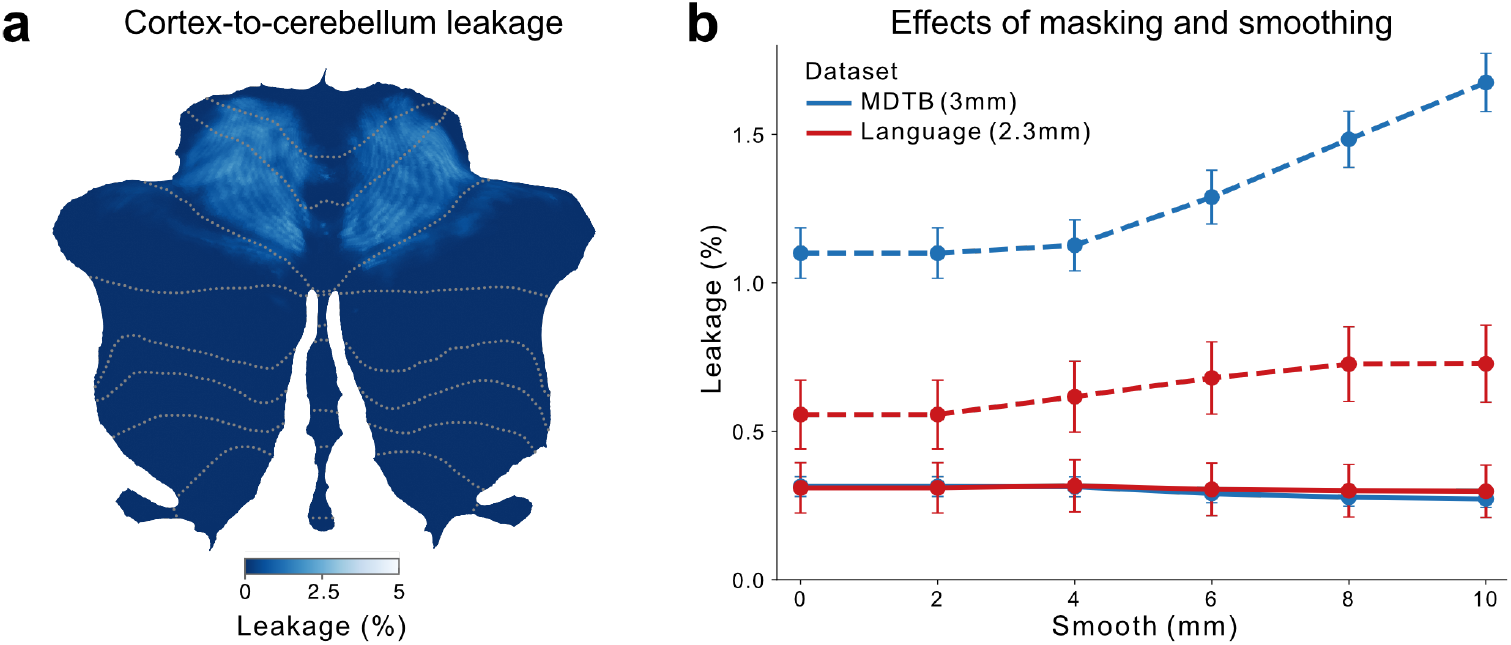
Cerebellar isolation mask reduces cortex-to-cerebellum signal leakage in fMRI analysis. **(a)** Cortex-to-cerebellum signal leakage visualized on the cerebellar flatmap. Because cortical tissue probability should be zero within the cerebellum, any non-zero cortical probability observed within cerebellar voxels reflects spurious signal leakage introduced during spatial reslicing and amplified by spatial smoothing. Shown is the group-averaged cortical probability map from the MDTB dataset after reslicing without cerebellar masking, at 6 mm FWHM smoothing. **(b)** Signal leakage quantified as the percentage of cortical signal within right lobule V, shown for simulations based on the anatomy and acquisition resolution of two functional datasets (MDTB, 3.0 mm; Language, 2.3 mm). Two processing strategies were compared: reslicing without masking (no mask, dashed lines), and standard reslicing with a U-Net-derived cerebellar mask (the default setting in SUITPy, solid lines). Error bars represent the standard error of the mean across subjects.

To prevent this leakage, we recommend masking of the functional maps with the cerebellar U-Net mask before reslicing the contrast maps into a cerebellar atlas space. This approach effectively suppressed cortical leakage across datasets and smoothing levels (Fig. 4b, solid lines). Importantly, subsequent smoothing within the cerebellum did not increase the amount of cortical signal. For most applications, the default functional analysis pipeline in the SUITPy toolbox therefore should suffice to prevent false-positives coming from neocortical activation.

The remaining cortical signal after masking comes from functional voxels that are located directly on the tentorium and therefore combine both supra-tentorial and sub-tentorial functional signals. The contribution of these voxels can be removed to zero by shrinking the U-Net mask further by excluding any voxels with non-zero neocortical grey and white matter probability.

## DISCUSSION

This paper presents SUITPy, a Python toolbox for cerebellar MRI analysis across the lifespan. The deep learning–based algorithm achieves robust and accurate cerebellar isolation across age groups and clinical populations, eliminating the need for manual correction. Cerebellum-only normalization consistently outperformed whole-brain approaches, with benefits extending across the full lifespan. Applying a cerebellar isolation mask in fMRI analysis effectively prevented contamination from adjacent neocortical signals, improving the specificity of cerebellar activation estimates. Beyond these core components, SUITPy provides a comprehensive analysis ecosystem, including support for multiple atlas spaces, a curated collection of functional and anatomical atlases for ROI-based analyses, and a 2D flatmap visualization of the cerebellar cortex.

### A new cerebellar isolation algorithm

For the Python version of SUIT, we developed a 3D U-Net based framework for cerebellar isolation from T1w/T2w MRI data. Trained on a diverse, manually corrected dataset spanning the full lifespan and including pathological cases, our model demonstrated robust and accurate performance across a wide range of anatomical variation. The U-Net consistently outperformed the probabilistic model implemented in previous Matlab-based versions of the SUIT toolbox (Diedrichsen, 2006). This advantage was reflected both in measures of boundary precision, as indicated by significantly lower Hausdorff distances (Huttenlocher et al., 1993), and in the standard Dice overlap measure.

Most importantly, the new U-Net based algorithm performed robustly on very young and old individuals, as well as on individuals with cerebellar degeneration, situations in which many of the existing algorithms fail. The quality of the resultant U-Net mask was good in general, such that manual correction is no longer necessary even for detailed anatomical analyses. Indeed, we showed that the quality of the cerebellum-only normalization was nearly equivalent whether a fully manual isolation mask or an automatically U-Net isolation mask was used. This makes the cerebellum-only pipeline easy to scale to large population-based datasets (Kim et al., 2025a,b; Moberget et al., 2018).

A key strength of our model is its ability to handle both T1w and T2w images with comparable accuracy, and to integrate both modalities for improved performance when available. This flexibility is particularly valuable in clinical and research settings where acquisition protocols vary or where one modality may be missing due to time constraints, motion artifacts, or retrospective data pooling. The practical benefit of incorporating a second modality is modest overall, and the cost-benefit trade-off of dual-modality acquisition should be considered in the context of the target population and available resources. The model’s success with T2w-only inputs also extends its utility to pediatric and fetal imaging, where T2w contrast often provides superior tissue differentiation.

The advantage of U-Net over our legacy probabilistic approach for cerebellar isolation shown here is in line with many other papers showing the power of this deep-learning approach across many other image segmentation problems (Faber et al., 2022; Çiçek et al., 2016; Li et al., 2018). The performance of any supervised learning model depends critically on the quality of the training data. Our manual isolation dataset, including 131 scans across eight distinct studies, represents a substantial investment in expert manual annotation. The resulting dataset, which spans the full lifespan and includes pathological cases, serves as a valuable resource for the neuroimaging community.

Despite its strengths, this study has several limitations that merit consideration. First, while our dataset encompassed substantial anatomical diversity, it remained limited in sample size (N=131) and did not capture the full spectrum of cerebellar pathologies. Rare congenital malformations, extreme atrophy, or post-surgical anatomies may fall outside the model’s training distribution and could produce less reliable predictions. Additionally, the model’s capacity on other clinical imaging modalities (e.g., FLAIR) remains to be tested.

### Cerebellar isolation mask improves subsequent spatial normalization

Our results demonstrated that spatial normalization of the cerebellum can be improved by masking the scan and normalizing it to a cerebellum-only template. This procedure improved registration accuracy and lobular alignment across individuals, even when the anatomical details in the cerebellar template and the normalization algorithm were the same as for whole-brain alignment. The benefits of cerebellum-only normalization are likely analogous to those of skull-stripping, which also improves subsequent spatial normalization by removing irrelevant surrounding tissue (Fischmeister et al., 2013; Ragguett et al., 2024). The masking step removes distracting and variable anatomical details from the image, so that the normalization algorithm can optimize the alignment of cerebellar structures specifically. Another benefit appears to be that the cerebellum-only normalization also works more robustly than whole-brain normalization of very young brains, where the comparable whole-brain pipeline often failed.

Normalization performance was reduced in neonatal datasets, consistent with the known inversion of gray–white matter contrast and rapid morphological changes during early postnatal development. However, even under these challenging conditions, the new U-Net–based cerebellar isolation algorithm performed well (Fig. 3f). Moreover, robust performance in infant datasets beyond the first postnatal year further supports the generalizability of the method (Fig. 3e). Nevertheless, the normalization performance can likely be further improved by using age-specific templates that reflect the tissue contrast in this age group (Hernandez-Castillo et al., 2019). Furthermore, incorporating both T1- and T2-weighted scans in the normalization step would likely improve estimates of the boundary between white and gray matter. We are currently refining the neonatal normalization framework based on these principles and will incorporate it into a future release of SUITPy to facilitate its broader use in cerebellar research.

The original version of the toolbox contained a specific cerebellar template, the SUIT template. Even though this template was generated to be spatially unbiased with respect to the affine MNI template (Diedrichsen, 2006), some structures differ in location by a few mm from the now standard non-linear MNI templates. The new toolbox, therefore, comes with two additional cerebellum-only templates: the `MNI152NLin6AsymC` and the `MNI152NLin2009cSymC`, which are masked versions of the `MNI152 NLin6Asym` and `MNI152NLin2009cSym` templates (Fonov et al., 2009). These templates allow the user to reap the benefits of a cerebellum-only pipeline without changing the atlas space for other brain structures. Of course, our results also show that the whole-brain pipeline using modern non-linear normalization algorithms and templates delivers relatively accurate results. However, in the new toolbox implementation, the isolation algorithm takes *<*1 minute per individual on a standard desktop or laptop computer without requiring a GPU. The subsequent normalization step takes only an additional 0.5–1 minute, such that the overall pipeline can be run very efficiently.

### Cerebellar isolation mask prevents bias from neocortical signals

Even when not used in the spatial normalization step, the cerebellar isolation mask should be used in fMRI analysis to reduce signal leakage from the inferior occipital cortex into cerebellar voxels. The issue is particularly important as the task-related activity in the neocortex is often much larger than in the cerebellum. Without masking, spatial smoothing of functional data, as is often done for fMRI group analysis, can therefore induce bias in lobules IV, V, and VI of the cerebellum.

Even with the standard masking applied in the toolbox, this leakage is not fully eliminated. This limitation reflects the partial volume effects across the tentorium. This remaining leakage can be reduced to zero by shrinking the U-Net mask further by excluding cortical voxels. Because this step will also reduce some real cerebellar activation, we generally recommend it only when there is a good reason to believe that the cerebellar activation could be artifactual – for example, when the functional activation found in the superior cerebellum is highly related to that in the directly abutting functional regions.

### A new Python-based SUIT toolbox

With the release `v2.1.0`, the isolation and normalization pipelines are implemented and extensively documented. The toolbox (Fig. 5) additionally provides a range of other functionalities. The newest version supports three cerebellum-only atlas spaces: the original `SUIT` template, and a cerebellar version of the common `MNI152NLin6Asym` template. Additionally, the symmetric `MNI152NLin2009cSym` template is especially useful for analyses of functional asymmetries, as ROIs can be defined easily in an equivalent fashion across left and right hemispheres (Nettekoven et al., 2024).

**Figure 5.**
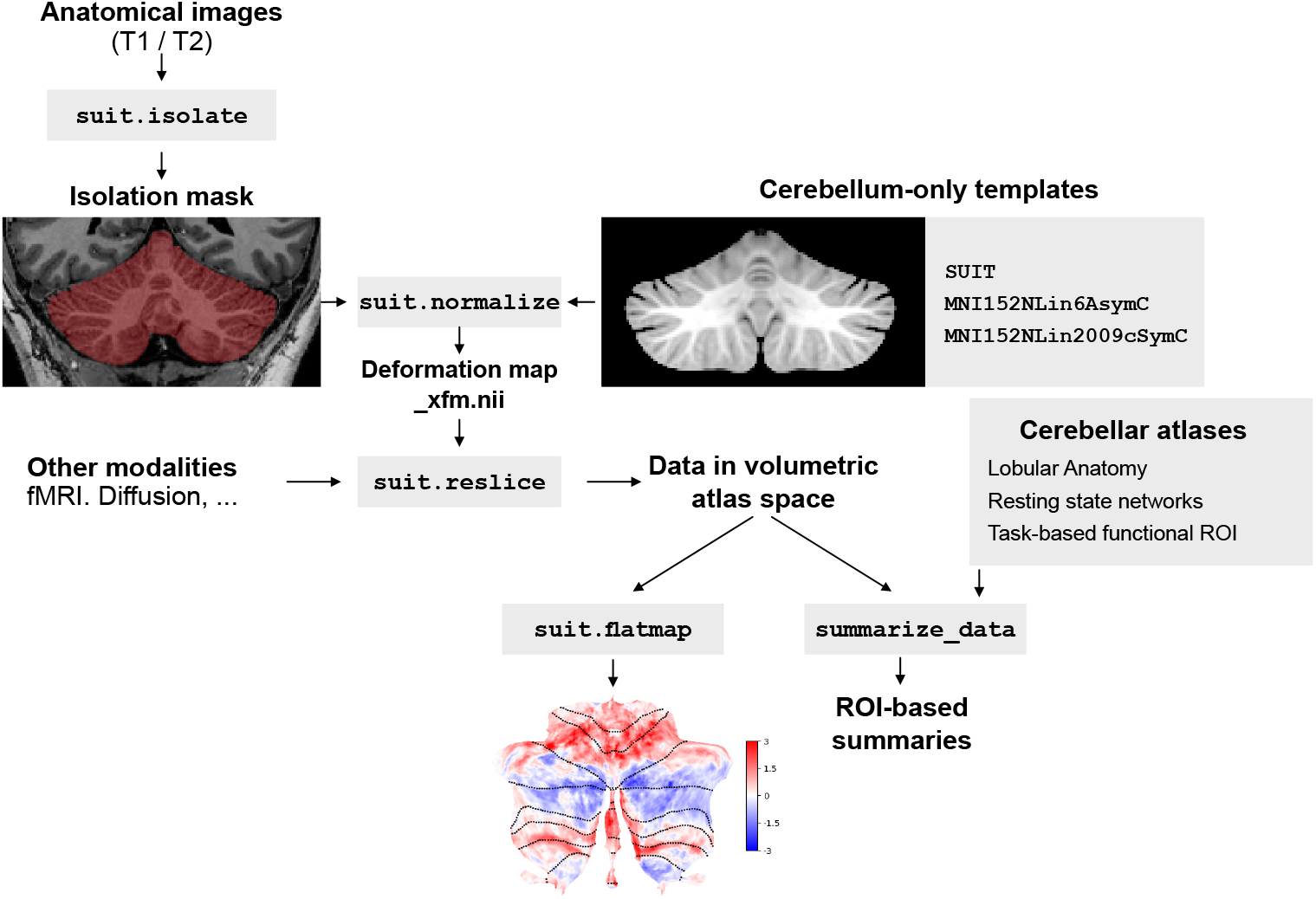
Functionality provided in the SUITPy toolbox. Apart from the described cerebellar isolation and cerebellum-only normalization, the toolbox provides functionality to reslice functional data into atlas space (after masking). The new toolbox supports three different cerebellum-only atlas spaces. Data in these spaces can be summarized using a wide range of anatomical and functional atlases, and can be visualized on a 2D-flatmap representation (Diedrichsen and Zotow, 2015).

The toolbox is associated with a collection of functional and anatomical atlases, including a probabilistic lobular atlas (Diedrichsen et al., 2009), atlases based on resting-state functional connectivity (Buckner et al., 2011; Ji et al., 2019), and task-based parcellations of the cerebellum (King et al., 2019; Nettekoven et al., 2024). These atlases can be used to provide ROI-based summaries of cerebellar data and to help interpret cerebellar findings based on the exact location. The repository also contains a collection of task-activity and feature maps. The toolbox is designed to be extensible, and community contributions are encouraged.

Another well-used feature of the toolbox is a 2D visualization of the cerebellar cortex (Diedrichsen and Zotow, 2015). Note that this flatmap does not attempt to unfold the actual highly convoluted individual cerebellar anatomy (Sereno et al., 2020), but rather to provide a 2D visualization that is at the appropriate scale for volume-based group averaged data that allows scientists to observe the functional activation state of the entire cerebellar grey matter in a single view. By flattening the complex folds of the cerebellum, the map reveals both the anterior-posterior (lobular) and medial-lateral structural organization, ensuring that localized activation clusters in the 3D volume remain visually intact on the 2D surface. Furthermore, the map is designed to minimize distortions, such that the surface area on the flatmap is roughly proportional to the represented gray-matter volume. The visualization is now widely used by the cerebellar research community.

In sum, SUITPy provides a voxel-wise pipeline for the human cerebellum that works robustly across the lifespan. It therefore provides an important complement to lobular segmentation approaches such as ACAPULCO (Han et al., 2020) and CerebNet (Faber et al., 2022). For fMRI studies, it improves group-based alignment and prevents signal leakage arising from abutting cortical structures. For anatomical studies, the improved normalization may improve the accuracy of voxel-wise normative modeling approaches (Chavanne et al., 2026) that aim to quantify inter-individual variability across the lifespan. SUITPy thus enables robust and accurate cerebellar mapping at scale, supporting population-level studies of cerebellar organization, development, and disease.

## ENDING SECTIONS

### Code Availability

The `SUITPy` toolbox (version 2.1) can be installed via `pip` with source code available at https://github.com/DiedrichsenLab/SUITPy. Tutorials and documentation can be found at https://suitpy.readthedocs.io.

### Dataset Availability

The manual isolation dataset (except for the EssCer dataset) is available upon request.

### Author Contributions

Conception: JD; Analysis: YL, YW, CHC; Implementation: YW, YL, BA, CN, VA, CHC, JD; Funding acquisition: AM, JD; Draft: YW, YL, JD; Final editing: all authors.

### Funding

The development of the new version of the toolbox was supported by a grant from the Raynor Cerebellar Project to JD. Additional support came from the Canadian Institutes of Health Research (CIHR, PJT-191815 to JD) and the Canada First Research Excellence Fund (BrainsCAN) to Western University. CN was supported by a Wellcome Trust Early Career Award (306553/Z/23/Z) and a Junior Research Fellowship Grant from Linacre College, University of Oxford.

## Acknowledgments

The authors thank Maedbh King, Da Zhi, Medha Porwal, Ladan Shahshahani, Julian Rochacz, and Matea Skenderija for technical help and feedback.

---

**Figure S1.**
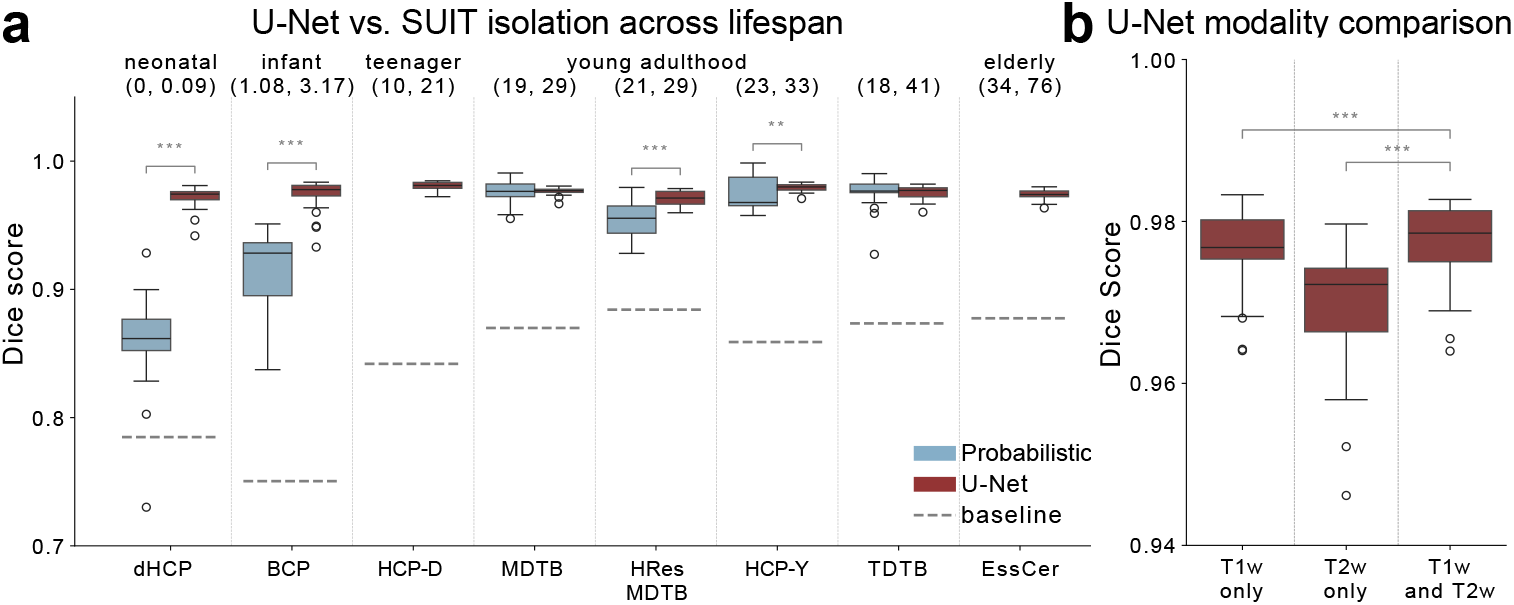
Dice score comparison of U-Net and probabilistic isolation algorithm. **(a)** Dice score for Probabilistic (blue) and U-Net (red) isolation. Dotted lines indicate baseline performance of the simple average of the training set. Horizontal bars with asterisks denote significant pairwise differences (paired t-tests, *: p<0.05, **: p<0.01, ***: p<0.001). **b**. Dice score comparison of U-Net isolation algorithm using different input modality configurations.

## REFERENCES

Arafat, B., Nettekoven, C., Xiang, J. D., and Diedrichsen, J. (2026). Multi-task batteries for precision functional mapping. bioRxiv.

Ashburner, J. (2007). A fast diffeomorphic image registration algorithm. Neuroimage, 38(1):95–113.

Ashburner, J. and Friston, K. J. (1999). Nonlinear spatial normalization using basis functions. Hum. Brain Mapp., 7(4):254–266.

Ashburner, J. and Friston, K. J. (2005). Unified segmentation. Neuroimage, 26(3):839–851.

Avants, B. B., Tustison, N. J., Song, G., Cook, P. A., Klein, A., and Gee, J. C. (2011). A reproducible evaluation of ANTs similarity metric performance in brain image registration. NeuroImage, 54(3):2033– 2044.

Baumann, O. and Mattingley, J. B. (2012). Functional topography of primary emotion processing in the human cerebellum. Neuroimage, 61(4):805–811.

Buckner, R. L., Krienen, F. M., Castellanos, A., Diaz, J. C., and Yeo, B. T. (2011). The organization of the human cerebellum estimated by intrinsic functional connectivity. J. Neurophysiol., 106(5):2322–2345.

Chavanne, A. V., Wang, Y., de Boer, A. A. A., Xu, B., van Prooije, T. H., Kapteijns, K. C. J., Reniers, C., Hernandez-Castillo, C. R., Fernandez-Ruiz, J., van de Warrenburg, B. P., Diedrichsen, J., Muetzel, R. L., and Marquand, A. F. (2026). Dissecting heterogeneous brain development and aging using voxelwise normative models. bioRxiv.

Chenarlogh, V. A., Hassanpour, A., Grolinger, K., and Parsa, V. (2024). Performance analysis of dilated one-to-many u-net model for medical image segmentation. IEEE Access, 12:197259–197274.

Çiçek, Ö., Abdulkadir, A., Lienkamp, S. S., Brox, T., and Ronneberger, O. (2016). 3d u-net: Learning dense volumetric segmentation from sparse annotation. In Ourselin, S., Joskowicz, L., Sabuncu, M. R., Unal, G., and Wells, W., editors, Medical Image Computing and Computer-Assisted Intervention – MICCAI 2016, pages 424–432, Cham. Springer International Publishing.

Cox, R. W. (1996). AFNI: Software for Analysis and Visualization of Functional Magnetic Resonance Neuroimages. Computers and Biomedical Research, 29(3):162–173.

Crum, W. R., Camara, O., and Hill, D. L. (2006). Generalized overlap measures for evaluation and validation in medical image analysis. IEEE transactions on medical imaging, 25(11):1451–1461.

Dice, L. R. (1945). Measures of the amount of ecologic association between species. Ecology, 26(3):297– 302.

Diedrichsen, J. (2006). A spatially unbiased atlas template of the human cerebellum. NeuroImage, 33(1):127–138.

Diedrichsen, J., Balsters, J. H. H., Flavell, J., Cussans, E., and Ramnani, N. (2009). A probabilistic MR atlas of the human cerebellum. Neuroimage, 46(1):39–46.

Diedrichsen, J. and Zotow, E. (2015). Surface-based display of volume-averaged cerebellar imaging data. PLoS One, 10(7):e0133402.

Draganova, R., Konietschke, F., Steiner, K. M., Elangovan, N., Gümüs, M., Göricke, S. M., Ernst, T. M., Deistung, A., Van Eimeren, T., Konczak, J., and Timmann, D. (2022). Motor training-related brain reorganization in patients with cerebellar degeneration. Human Brain Mapping, 43(5):1611–1629.

Edwards, A. D., Rueckert, D., Smith, S. M., Abo Seada, S., Alansary, A., Almalbis, J., Allsop, J., Andersson, J., Arichi, T., Arulkumaran, S., Bastiani, M., Batalle, D., Baxter, L., Bozek, J., Braithwaite, E., Brandon, J., Carney, O., Chew, A., Christiaens, D., Chung, R., Colford, K., Cordero-Grande, L., Counsell, S. J., Cullen, H., Cupitt, J., Curtis, C., Davidson, A., Deprez, M., Dillon, L., Dimitrakopoulou, K., Dimitrova, R., Duff, E., Falconer, S., Farahibozorg, S.-R., Fitzgibbon, S. P., Gao, J., Gaspar, A., Harper, N., Harrison, S. J., Hughes, E. J., Hutter, J., Jenkinson, M., Jbabdi, S., Jones, E., Karolis, V., Kyriakopoulou, V., Lenz, G., Makropoulos, A., Malik, S., Mason, L., Mortari, F., Nosarti, C., Nunes, R. G., O’Keeffe, C., O’Muircheartaigh, J., Patel, H., Passerat-Palmbach, J., Pietsch, M., Price, A. N., Robinson, E. C., Rutherford, M. A., Schuh, A., Sotiropoulos, S., Steinweg, J., Teixeira, R. P. A. G., Tenev, T., Tournier, J.-D., Tusor, N., Uus, A., Vecchiato, K., Williams, L. Z. J., Wright, R., Wurie, J., and Hajnal, J. V. (2022). The Developing Human Connectome Project Neonatal Data Release. Frontiers in neuroscience, 16:886772. Place: Switzerland.

Faber, J., Kügler, D., Bahrami, E., Heinz, L.-S., Timmann, D., Ernst, T. M., Deike-Hofmann, K., Klockgether, T., van de Warrenburg, B., van Gaalen, J., Reetz, K., Romanzetti, S., Oz, G., Joers, J. M., Diedrichsen, J., and Reuter, M. (2022). CerebNet: A fast and reliable deep-learning pipeline for detailed cerebellum sub-segmentation. NeuroImage, 264:119703. Place: United States.

Fischmeister, F. P. S., Höllinger, I., Klinger, N., Geissler, A., Wurnig, M. C., Matt, E., Rath, J., Robinson, S. D., Trattnig, S., and Beisteiner, R. (2013). The benefits of skull stripping in the normalization of clinical fMRI data. NeuroImage Clin., 3:369–380.

Fonov, V., Evans, A. C., Botteron, K., Almli, C. R., McKinstry, R. C., and Collins, D. L. (2011). Unbiased average age-appropriate atlases for pediatric studies. NeuroImage, 54(1):313–327.

Fonov, V. S., Evans, A. C., McKinstry, R. C., Almli, C. R., and Collins, D. (2009). Unbiased nonlinear average age-appropriate brain templates from birth to adulthood. NeuroImage, 47:S102.

Grabner, G., Janke, A. L., Budge, M. M., Smith, D., Pruessner, J., and Collins, D. L. (2006). Symmetric atlasing and model based segmentation: An application to the hippocampus in older adults. In Larsen, R., Nielsen, M., and Sporring, J., editors, Medical Image Computing and Computer-Assisted Intervention – MICCAI 2006, pages 58–66, Berlin, Heidelberg. Springer Berlin Heidelberg.

Han, S., Carass, A., He, Y., and Prince, J. L. (2020). Automatic cerebellum anatomical parcellation using U-Net with locally constrained optimization. NeuroImage, 218:116819. Place: United States.

Hatamizadeh, A., Tang, Y., Nath, V., Yang, D., Myronenko, A., Landman, B., Roth, H. R., and Xu, D. (2022). Unetr: Transformers for 3d medical image segmentation. In Proceedings of the IEEE/CVF winter conference on applications of computer vision, pages 574–584.

Hernandez-Castillo, C. R., Limperopoulos, C., and Diedrichsen, J. (2019). A representative template of the neonatal cerebellum. Neuroimage, 184.

Howell, B. R., Styner, M. A., Gao, W., Yap, P.-T., Wang, L., Baluyot, K., Yacoub, E., Chen, G., Potts, T., Salzwedel, A., Li, G., Gilmore, J. H., Piven, J., Smith, J. K., Shen, D., Ugurbil, K., Zhu, H., Lin, W., and Elison, J. T. (2019). The UNC/UMN Baby Connectome Project (BCP): An overview of the study design and protocol development. NeuroImage, 185:891–905. Place: United States.

Huang, H., Lin, L., Tong, R., Hu, H., Zhang, Q., Iwamoto, Y., Han, X., Chen, Y.-W., and Wu, J. (2020). Unet 3+: A full-scale connected unet for medical image segmentation. In ICASSP 2020 - 2020 IEEE International Conference on Acoustics, Speech and Signal Processing (ICASSP), pages 1055–1059.

Huttenlocher, D., Klanderman, G., and Rucklidge, W. (1993). Comparing images using the hausdorff distance. IEEE Transactions on Pattern Analysis and Machine Intelligence, 15(9):850–863.

Isensee, F., Petersen, J., Klein, A., Zimmerer, D., Jaeger, P. F., Kohl, S., Wasserthal, J., Koehler, G., Norajitra, T., Wirkert, S., and Maier-Hein, K. H. (2018). nnu-net: Self-adapting framework for u-net-based medical image segmentation.

Ji, J. L., Spronk, M., Kulkarni, K., Repovš, G., Anticevic, A., and Cole, M. W. (2019). Mapping the human brain’s cortical-subcortical functional network organization. Neuroimage, 185:35–57.

Kim, M., Leonardsen, E. H., Rutherford, S., Selbæk, G., Persson, K., Steen, N. E., Smeland, O. B., Richard, G., Alnæs, D., Boen, R., Sønderby, I. E., van der Meer, D., Andreassen, O. A., Westlye, L. T., Wolfers, T., and Moberget, T. (2025a). Mapping cerebellar morphology in 15q11.2 CNV carriers using normative modeling. medRxiv.

Kim, M., Sharma, N., Leonardsen, E. H., Rutherford, S., Selbæk, G., Persson, K., Steen, N. E., Smeland, O. B., Ueland, T., Richard, G., Manoli, A., Valk, S. L., Alnæs, D., Beckman, C. F., Marquand, A. F., Andreassen, O. A., Westlye, L. T., Wolfers, T., and Moberget, T. (2025b). Predicting mental and neurological illnesses based on cerebellar normative features. Biol. Psychiatry Glob. Open Sci., 5(5):100541.

King, M., Hernandez-Castillo, C. R., Poldrack, R. A., Ivry, R. B., and Diedrichsen, J. (2019). Functional boundaries in the human cerebellum revealed by a multi-domain task battery. Nature Neuroscience, 22(8):1371–1378.

Kingma, D. P. and Ba, J. (2017). Adam: A method for stochastic optimization.

Leiner, H., Leiner, A., and Dow, R. (1986). Does the cerebellum contribute to mental skills? Behavioral Neuroscience, 100:443–454.

Li, X., Chen, H., Qi, X., Dou, Q., Fu, C.-W., and Heng, P.-A. (2018). H-denseunet: Hybrid densely connected unet for liver and tumor segmentation from ct volumes. IEEE Transactions on Medical Imaging, 37(12):2663–2674.

Maas, A. L. (2013). Rectifier nonlinearities improve neural network acoustic models. McCarthy, P. (2025). FSLeyes.

Moberget, T., Doan, N. T., Alnæs, D., Kaufmann, T., Córdova-Palomera, A., Lagerberg, T. V., Diedrichsen, J., Schwarz, E., Zink, M., Eisenacher, S., Kirsch, P., Jönsson, E. G., Fatouros-Bergman, H., Flyckt, L., Pergola, G., Quarto, T., Bertolino, A., Barch, D., Meyer-Lindenberg, A., Agartz, I., Andreassen, O. A., and Westlye, L. T. (2018). Cerebellar volume and cerebellocerebral structural covariance in schizophrenia: a multisite mega-analysis of 983 patients and 1349 healthy controls. Mol. Psychiatry, 23(6):1512–1520.

Nettekoven, C., Zhi, D., Shahshahani, L., Pinho, A. L., Saadon-Grosman, N., Buckner, R. L., and Diedrichsen, J. (2024). A hierarchical atlas of the human cerebellum for functional precision mapping. Nature Communications, 15(1):8376.

Phillips, J. R., Hewedi, D. H., Eissa, A. M., and Moustafa, A. A. (2015). The cerebellum and psychiatric disorders. Frontiers in public health, 3:66. Place: Switzerland.

Ragguett, R.-M., Eagleson, R., and de Ribaupierre, S. (2024). Evaluating normalized registration and preprocessing methodologies for the analysis of brain MRI in pediatric patients with shunt-treated hydrocephalus. Front. Neurosci., 18:1405363.

Ronneberger, O., Fischer, P., and Brox, T. (2015). U-Net: Convolutional Networks for Biomedical Image Segmentation. In Navab, N., Hornegger, J., Wells, W. M., and Frangi, A. F., editors, Medical Image Computing and Computer-Assisted Intervention – MICCAI 2015, pages 234–241, Cham. Springer International Publishing.

Sayın, K. A., Gürsoy, N. K., Yolcu, T., and Gürsoy, A. (2025). On the synergy of optimizers and activation functions: A cnn benchmarking study. Mathematics, 13(13):2088.

Schmahmann, J. D. (2004). Disorders of the Cerebellum: Ataxia, Dysmetria of Thought, and the Cerebellar Cognitive Affective Syndrome. The Journal of Neuropsychiatry and Clinical Neurosciences, 16:367–378.

Sereno, M. I., Diedrichsen, J., Tachrount, M., Testa-Silva, G., d’Arceuil, H., and De Zeeuw, C. (2020). The human cerebellum has almost 80% of the surface area of the neocortex. Proceedings of the National Academy of Sciences, 117(32):19538–19543.

Shahshahani, L., King, M., Nettekoven, C., Ivry, R. B., and Diedrichsen, J. (2024). Selective recruitment of the cerebellum evidenced by task-dependent gating of inputs. eLife, 13:1–19.

Somerville, L. H., Bookheimer, S. Y., Buckner, R. L., Burgess, G. C., Curtiss, S. W., Dapretto, M., Elam, J. S., Gaffrey, M. S., Harms, M. P., Hodge, C., Kandala, S., Kastman, E. K., Nichols, T. E., Schlaggar, B. L., Smith, S. M., Thomas, K. M., Yacoub, E., Van Essen, D. C., and Barch, D. M. (2018). The Lifespan Human Connectome Project in Development: A large-scale study of brain connectivity development in 5-21 year olds. NeuroImage, 183:456–468. Place: United States.

Strick, P. L., Dum, R. P., and Fiez, J. A. (2009). Cerebellum and nonmotor function. Annual review of neuroscience, 32:413–434. Place: United States.

Studholme, C., Hill, D., and Hawkes, D. (1999). An overlap invariant entropy measure of 3d medical image alignment. Pattern Recognition, 32(1):71–86.

Sørensen, T. (1948). A method of establishing groups of equal amplitude in plant sociology based on similarity of species and its application to analyses of the vegetation on danish commons.

Timmann, D., Brandauer, B., Hermsdörfer, J., Ilg, W., Konczak, J., Gerwig, M., Gizewski, E. R., and Schoch, B. (2008). Lesion-symptom mapping of the human cerebellum. Cerebellum, 7(4):602–606.

Tustison, N. J., Avants, B. B., Cook, P. A., Yuanjie Zheng, Egan, A., Yushkevich, P. A., and Gee, J. C. (2010). N4ITK: Improved N3 Bias Correction. IEEE Transactions on Medical Imaging, 29(6):1310– 1320.

Ulyanov, D., Vedaldi, A., and Lempitsky, V. (2016). Instance normalization: The missing ingredient for fast stylization. arXiv preprint arXiv:1607.08022.

Van Essen, D. C., Ugurbil, K., Auerbach, E., Barch, D., Behrens, T. E. J., Bucholz, R., Chang, A., Chen, L., Corbetta, M., Curtiss, S. W., Della Penna, S., Feinberg, D., Glasser, M. F., Harel, N., Heath, A. C., Larson-Prior, L., Marcus, D., Michalareas, G., Moeller, S., Oostenveld, R., Petersen, S. E., Prior, F., Schlaggar, B. L., Smith, S. M., Snyder, A. Z., Xu, J., and Yacoub, E. (2012). The Human Connectome Project: a data acquisition perspective. NeuroImage, 62(4):2222–2231. Place: United States.

Zhou, Z., Rahman Siddiquee, M. M., Tajbakhsh, N., and Liang, J. (2018). Unet++: A nested u-net architecture for medical image segmentation. In Stoyanov, D., Taylor, Z., Carneiro, G., Syeda-Mahmood, T., Martel, A., Maier-Hein, L., Tavares, J. M. R., Bradley, A., Papa, J. P., Belagiannis, V., Nascimento, J. C., Lu, Z., Conjeti, S., Moradi, M., Greenspan, H., and Madabhushi, A., editors, Deep Learning in Medical Image Analysis and Multimodal Learning for Clinical Decision Support, pages 3–11, Cham. Springer International Publishing.

